# Cell-to-cell heterogeneities during extrinsic apoptosis arise from cell cycle progression and transmitotic apoptosis resistance

**DOI:** 10.1101/2021.02.26.433034

**Authors:** Nadine Pollak, Aline Lindner, Dirke Imig, Karsten Kuritz, Jacques S. Fritze, Isabel Heinrich, Jannis Stadager, Stephan Eisler, Daniela Stöhr, Frank Allgöwer, Peter Scheurich, Markus Rehm

## Abstract

Extrinsic apoptosis relies on TNF-family receptor activation by immune cells or receptor-activating biologics. Here, we monitored cell cycle progression at minutes resolution to relate apoptosis kinetics and cell-to-cell heterogeneities in death decisions to cell cycle phases. Interestingly, we found that cells in S phase delay TRAIL receptor-induced death in favour for mitosis, thereby passing on an apoptosis-primed state to their offspring. This translates into two distinct fates, apoptosis execution post mitosis or cell survival from inefficient apoptosis. Transmitotic resistance is linked to Mcl-1 upregulation from mid S phase onwards, which allows cells to pass through mitosis with activated caspase-8, and with cells escaping apoptosis after mitosis sustaining sublethal DNA damage. Antagonizing Mcl-1 by BH3-mimetics suppresses cell cycle-dependent delays in apoptosis, prevents apoptosis-resistant progression through mitosis and averts unwanted survival from apoptosis induction. Cell cycle progression therefore modulates signal transduction during extrinsic apoptosis, with Mcl-1 governing decision making between death, proliferation and survival from inefficient apoptosis induction. Cell cycle progression thus is a crucial process from which cell-to-cell heterogeneities in fates and treatment outcomes emerge in isogenic cell populations during extrinsic apoptosis signalling.

## Introduction

Healthy development and physiological organ functions rely on appropriately balanced cell proliferation and cell death, and consequently dysregulations in this balance manifest in both proliferative and degenerative diseases (1, 2). Proliferation signalling drives progression through the cell cycle and accumulation of cell mass, whereas apoptosis, the major form of programmed cell death, eliminates cells by triggering proteolytic cascades in which caspases act as cell death initiators and executors. Two main apoptosis pathways activate caspases. The extrinsic pathway is triggered by death ligands binding to their respective cell surface receptor(s), whereas the intrinsic, mitochondrial pathway is initiated in response to cytotoxic stress (3). The extrinsic pathway can be triggered by immune cells to induce apoptosis in transformed and potentially harmful cells (4), and death ligand TRAIL now serves as a template for the development of novel anti-cancer therapeutics and treatment strategies, with 2^nd^ generation ligands recently having entered clinical trials (5).

Following ligand binding, activated death receptors cluster and form the death-inducing signalling complex (DISC), the activation platform for initiator caspases-8/-10. Active caspases-8/-10 in select cell types sufficiently cleave and activate effector caspase-3, resulting in type I apoptosis (6). Typically, however, the presence of the inactive caspase-8 homologue cFLIP as well as caspase-3 inhibitor XIAP prevent this type of apoptosis. Instead, the majority of cells rely on the amplification of the apoptotic signal via the mitochondrial pathway, i.e. type II signalling (7–9). Here, caspase-8 cleaves the BH3-only protein Bid, which then translocates to the mitochondria and stimulates Bax/Bak-dependent mitochondrial outer membrane permeabilisation (MOMP), a process that is inhibited by anti-apoptotic Bcl-2 family members such as Bcl-2, Bcl-xL and Mcl-1 (10). MOMP results in the release of cytochrome c and Smac from the mitochondria, driving efficient caspase-3 activation and apoptosis execution (11).

While prolonged cell cycle arrest, for example by cytotoxic drugs such as doxorubicin, CPT-11, bortezomib, simvastatin or anti-microtubule agent taxol, evidently induces stress responses that sensitize to extrinsic apoptosis (12–16), it so far remains unsolved if extrinsic apoptosis susceptibility is modulated throughout regular cell cycle progression. This may not be surprising, since combined and parallel monitoring of externally unperturbed cell cycle progression and apoptosis signalling is challenging. For example, bulk analyses of chemically synchronized cell populations provide little temporal resolution and suffer from interfering with normal cell cycle progression (17). Single cell time-lapse analyses, instead, so far remained restricted to studies conducted in the presence of apoptosis sensitizers, such as translation inhibitor cycloheximide (18, 19). Since cell cycle progression heavily relies on concerted protein production and degradation, and since short lived modulators of extrinsic apoptosis responsiveness, such as cFLIP, are rapidly lost upon translation inhibition (20), the question of whether cell cycle progression modulates extrinsic apoptosis signal transduction remains largely unaddressed so far.

Here, we therefore combined mathematical modelling and live-cell imaging to quantitatively study TRAIL-induced extrinsic apoptosis during unperturbed cell cycle progression. Interestingly, we found that cells progressing towards mitosis delay apoptotic signal transduction, attributable to an intrinsic, cell cycle associated upregulation of Mcl-1 and, to a lower extent, Bcl-xL. These cells transition through mitosis with activated caspase-8, frequently escape cell death, but show signs of sublethal DNA damage. Importantly, Mcl-1-targeting BH3-mimetics suffice to prevent unwanted transmitotic escape from extrinsic apoptosis.

## Results

### TRAIL-treated cells delay apoptosis execution in favour of mitosis

To analyse if cell cycle progression influences the timing of apoptosis execution in response to TRAIL treatment, we employed a combined approach of mathematical modelling and experimental observation of cell divisions and cell death events. To non-invasively monitor cell cycle phases, we generated NCI-H460 and HCT-116 cells stably expressing the fluorescently tagged degron of geminin (21, 22), which intrinsically labels cells throughout S, G_2_ and M phases, but not in G_1_ (**Figure S1a**). Treatment with a translationally relevant TRAIL receptor (TRAIL-R) agonist (Fc-scTRAIL) (23) dose-dependently reduced cell viability, accompanied by cleavage of caspase-8, caspase-3 as well as PARP (**Figure S1b, c**). Addition of pan-caspase inhibitor QVD-OPh averted cell death, demonstrating that cell death proceeded entirely by apoptosis (**Figure S1d**). Following TRAIL addition, we microscopically recorded if cells executed apoptosis prior to cell division or instead underwent mitosis before executing apoptosis. These data were then compared to expected patterns of apoptosis execution obtained from a mathematical cell population model in which progression towards mitosis and the timing of apoptosis were implemented as independent processes (**Figure 1a**) (24). While modelling predicted that approximately 20% of the NCI-H460 and HCT-116 cell populations would be expected to divide before executing apoptosis, significantly more cells underwent cell division prior to apoptosis in these experiments (**Figure 1b**). Closer examination of the imaging data indeed revealed that geminin-positive cells in S/G_2_ phases frequently first underwent mitosis, with daughter cells dying soon after cytokinesis (**Figure 1c, Figure S1e** and **Movie**). Correspondingly, when both model and experimental analyses were constrained to cells in S/G_2_/M phases at the time of TRAIL addition, the mismatch in the timing between apoptosis execution and mitosis was even more pronounced (**Figure 1d**). The comparison of mathematical predictions and experimental observations of cell death and cell division events therefore suggests that cell cycle progression might delay or decelerate the kinetics of apoptosis signal transduction, and that such delays would be expected to manifest predominantly during the S/G_2_/M phases of the cell cycle.

**Figure 1:**
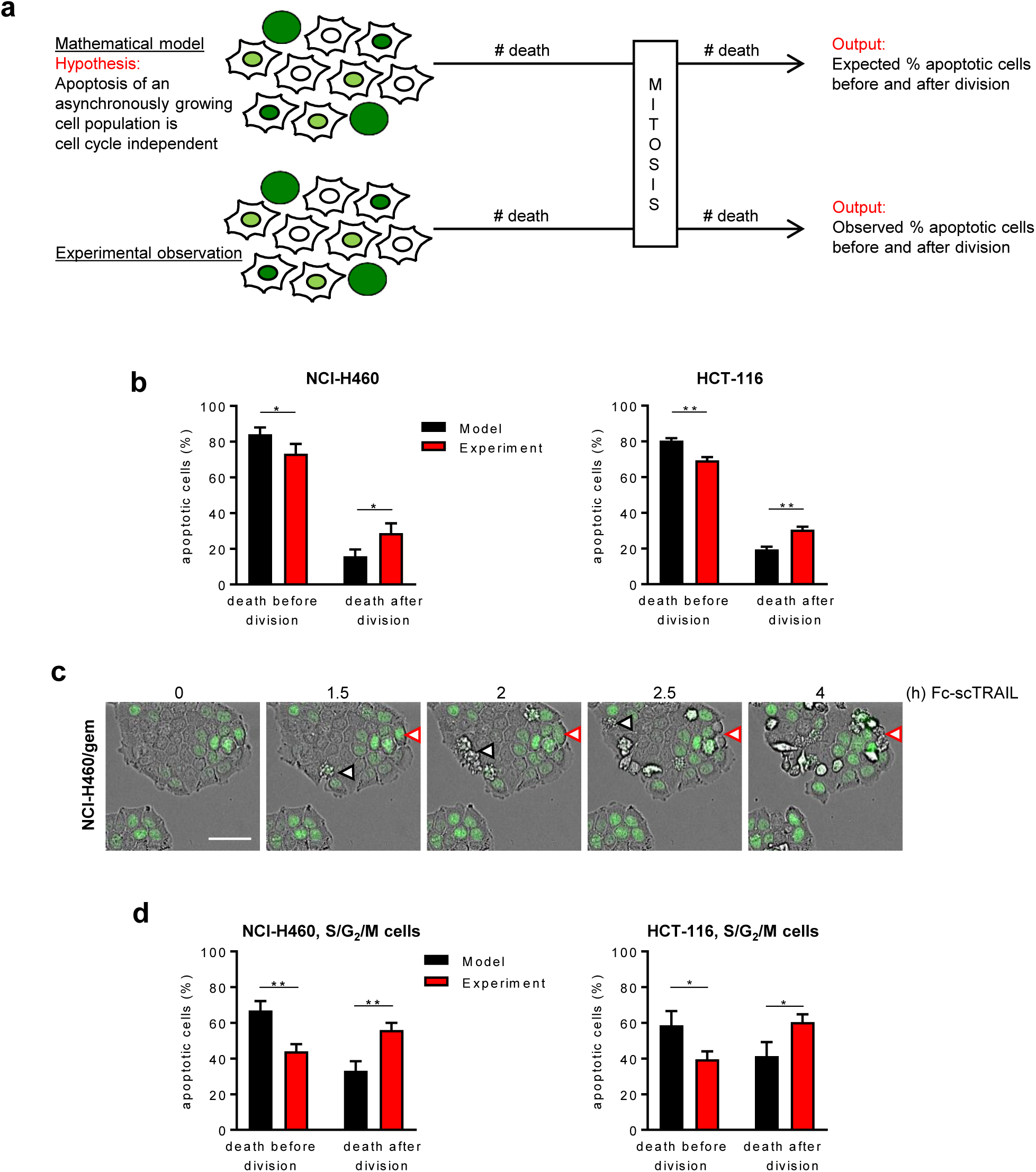
TRAIL-treated cells delay apoptosis execution in favour of mitosis. **a** Schematic representation of comparing mathematically modelled and experimentally observed cell cycle progression and apoptosis in response to TRAIL receptor stimulation. **b** Expected and observed cell death events in relation to cell division. Cells were treated with Fc-scTRAIL (0.06 nM) and monitored for 24 h. The percentage of apoptosis events before or after division was analysed for mathematically modelled or experimentally observed cell populations. Data are from at least 80-90 cells from three independent model runs or experiments and shown as mean + SD. *p<0.05, **p<0.01, unpaired t-tests. **c** Representative time-lapse images of NCI-H460/geminin cells treated with Fc-scTRAIL (0.06 nM). Scale bar = 50 µm. Cells dying prior to division are highlighted by black arrowheads. Red arrowhead tracks a cell dying after division. **d** Expected and observed cell death events of S/G_2_/M cells in relation to cell division. Analyses were performed as in (B). *p<0.05, **p<0.01, unpaired t-tests.

### Commitment to mitosis and delay of apoptosis are decided at mid to end of the S phase

We next experimentally studied if signal transduction kinetics towards apoptosis execution indeed differed between cells exposed to TRAIL at different times of the cell cycle and, if this is the case, at which point during cell cycle progression the decision is made to delay apoptosis in favour of mitosis. To this end, we monitored untreated cells for at least 20 h, so that cell divisions could be observed and the age of each individual cell was known at minutes resolution at the time of TRAIL addition. In parallel, we tracked the durations of cell cycle phases based on geminin intensity and cell morphology, as well as the times from TRAIL addition to apoptosis execution (**Figure 2a**). At population level, NCI-H460 and HCT-116 cells underwent apoptosis as soon as 1 h following TRAIL addition, with a median t_death_ of approximately 3 h (2.8 h and 3.7 h, respectively) (**Figure 2b, c**, orange). When separating cell populations based on the cell cycle phase in which they were exposed to TRAIL (**Figure 2a**), we noted that cells exposed to TRAIL in S/G_2_/M phases required significantly longer to die than cells exposed to TRAIL in G_1_ phase (**Figure 2b, c**). The delays observed in the S/G_2_/M populations could be ascribed to cells dying after mitosis (t_death_ calculated from f0 and f1 cell generations) (**Figure 2b, c**). Similar results were obtained in HeLa and HT1080 cells treated with TRAIL, in NCI-H460 cells treated with FasL as an alternative death ligand, and in two primary cell isolates obtained from melanoma metastases (**Figure S2a-d**).

**Figure 2:**
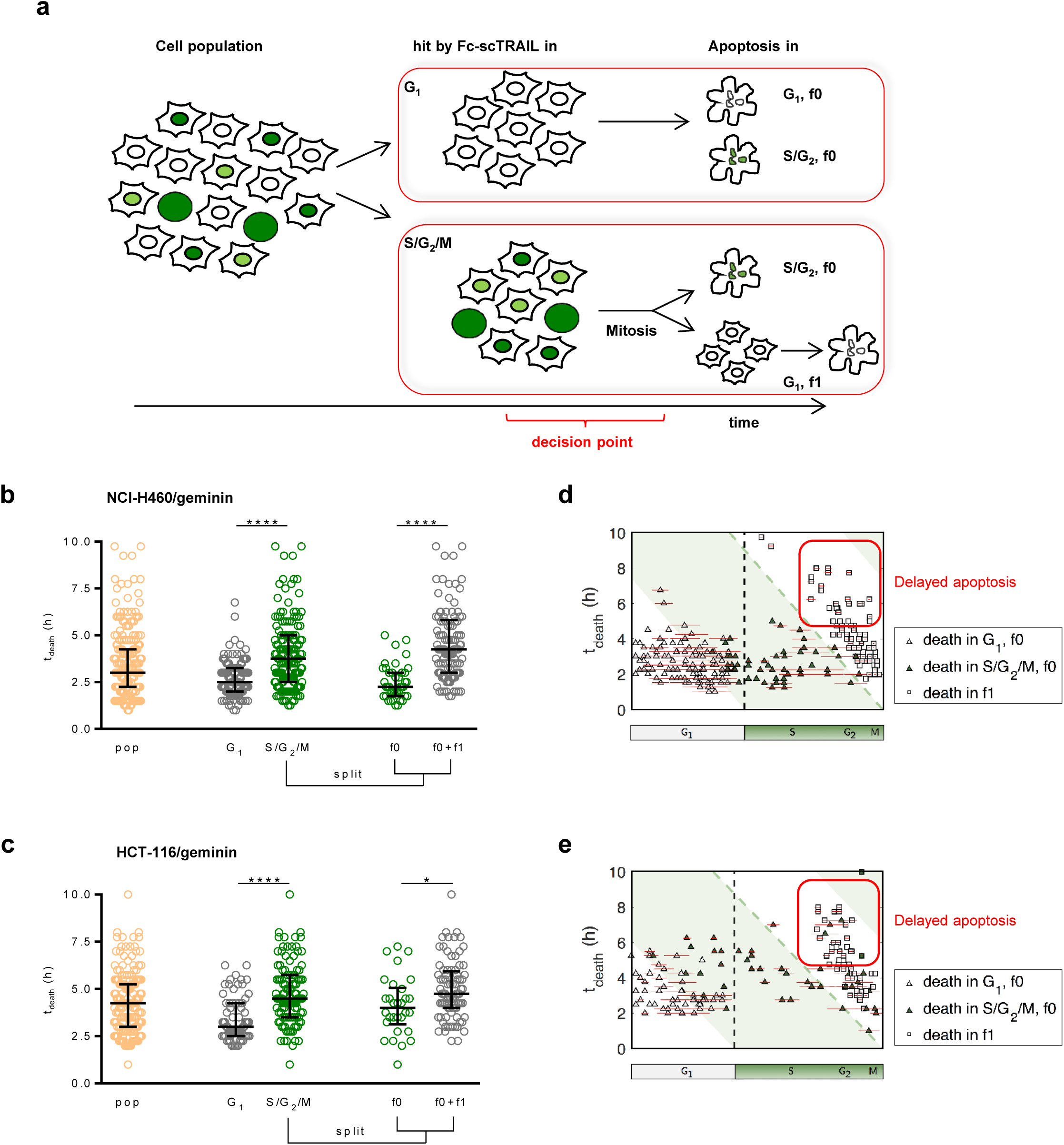
Commitment to mitosis and delay of apoptosis are decided at mid to end of the S phase. **a** Schematic representation of kinetic analyses. Unsynchronized cell populations were monitored for prolonged times and separated into G_1_ and S/G_2_/M subpopulations at the time of Fc-scTRAIL addition, based on geminin intensities. Times required to die (t_death_) were recorded for each cell, taking into account that mitosis might precede cell death. Subsequent analyses aimed at characterizing death kinetics and identifying the decision point at which cells preferentially undergo mitosis prior to executing apoptosis. **b and c**Times required to die (t_death_) of NCI-H460/geminin (B) or HCT-116/geminin (C) cells following treatment with Fc-scTRAIL (0.06 nM). Cells were monitored for 20 h prior to Fc-scTRAIL addition to ensure the age of each cell was known (time since previous division, mitotic history). Randomly chosen cells in different cell cycle stages were tracked and t_death_ of individual cells was plotted according the population separation described in (A). Plots show medians, 2^nd^ and 3^rd^ quartiles from at least 180 cells from at least three independent experiments (**** p<0.0001, * p<0.05 Mann-Whitney U test). **d and e** t_death_ times were plotted against cell cycle positions at the time of Fc-scTRAIL exposure. White areas represent G_1,_ green areas mark S/G_2_/M phases. Dashed green line denotes cell division. Triangles and squares represent individual cells, with red error bars indicating uncertainty estimates. n = 271 and n = 183 cells were analysed, respectively. Cells that progress through mitosis frequently show a significant delay in apoptosis execution (red boxes).

We next sought to identify when during the S/G_2_/M phases cells decide to delay apoptosis in favour of mitosis. Following data normalisation (24), we plotted the death times t_death_ of individual cells against their cell cycle position. This analysis revealed that cells notably delaying apoptosis execution following TRAIL exposure had completed approximately mid to late S phase (**Figure 2d, e**).

Taken together, these results demonstrate that many cells that would have had sufficient time to die prior to mitosis instead undergo cell division and delay extrinsic apoptosis, a behaviour that manifests once cells have entered mid to late S phase.

### Caspase-8 remains active throughout mitosis, but efficient caspase-3 activation is suppressed during G_2_/M phases

We next studied at which point during signal transduction the propagation of extrinsic apoptotic signalling is delayed. Surface amounts of TRAIL-R1/2 continuously increased from G_1_ throughout S/G_2_ and M phases in both NCI-H460 and HCT-116 cells, with M phase amounts approximately twice as high as in G_1_ (**Figure S2e, f**), as would be expected for proteins whose steady state amounts correlate with cell growth (25). TRAIL binding correlated with these changes (**Figure S2g, h**). These results therefore exclude that delays in apoptosis signalling emanate from lack of receptors or impaired ligand binding in later phases of the cell cycle. Next, we studied caspase-8 processing and activity in cells exposed to TRAIL at different times of the cell cycle. Since the activation of downstream caspases interferes with measurements of initiator caspase processing and activities (26), we used HCT-116 (Bax/Bak)^-/-^ cells (**Figure S2i**), incompetent to undergo MOMP and apoptosis execution, in our initial analyses. HCT-116 (Bax/Bak)^-/-^ cells, sorted by geminin fluorescence for G_1_ or G_2_/M phases, similarly processed procaspase-8 into p43/41 and p18 subunits (**Figure 3a**). We next studied if caspase-8 processing results in notable activity during S/G_2_/M phases. To this end, we introduced a FRET probe carrying the optimal recognition motif (IETD) for caspase-8 dependent proteolysis (27) into HCT-116 (Bax/Bak)^-/-^ or NCI-H460/Bcl-2 (**Figure S3a**) cells. Time-lapse imaging revealed caspase-8 activities in late stages of the cell cycle and that cells can enter and pass through mitosis with activated caspase-8 (**Figure 3b, S3b**). Corresponding to these observations, the cleavage of Bid, a natural substrate of caspase-8 that links caspase-8 activity to the mitochondrial apoptosis pathway and that is essential for TRAIL induced viability loss in NCI-H460 and HCT-116 cells (**Figure 3c**), was indistinguishable between cells in G_1_ or G_2_/M phases (**Figure 3d**). Caspase-8 additionally cleaves and activates effector procaspase-3. In cells relying on the mitochondrial apoptosis pathway, x-linked inhibitor of apoptosis protein (XIAP) suppresses full autoproteolytic maturation of the large caspase-3 subunit (8, 28). Consequently, in absence of MOMP and amplification of apoptosis signalling, caspase-3 processing is limited and only the initial cleavage product of the large caspase-3 subunit can be detected. Indeed, this was seen when comparing caspase-3 processing between HCT-116 (Bax/Bak)^-/-^ cells, parental HCT-116 cells and NCI-H460 cells (**Figure 3e**). Similar patterns of caspase-3 processing were observed in Bid depleted cells (**Figure 3f**). Staining HCT-116 (Bax/Bak)^-/-^ cells for DNA and processed caspase-3, followed by flow cytometric analysis, revealed identical procaspase-3 processing in G_1_, S and G_2_/M phases (**Figure 3gi**). In MOMP-competent HCT-116 cells, instead, additional peaks for cells with fully cleaved caspase-3 pools could be detected. Interestingly, these were restricted to cells in G_1_ and S phases, indicating that cells in G_2_/M phases suppress full processing of caspase-3 (**Figure 3gii**). NCI-H460 cells similarly limited caspase-3 processing in G_2_/M phases, indicative of evading the post-MOMP phase of apoptosis execution (**Figure 3giii**). When quantifying the kinetics of procaspase-3 cleavage events in the distinct subpopulations, we noted that initial procaspase-3 cleavage occurred cell cycle-independently in both NCI-H460 and HCT-116 cells (**Figure 3h**). Pre- and post-MOMP caspase-3 processing followed very similar kinetics in cells from G_1_ and S phases, whereas the post-MOMP subpopulation remained very low in cells analysed in G_2_/M phases (**Figure 3h**).

**Figure 3:**
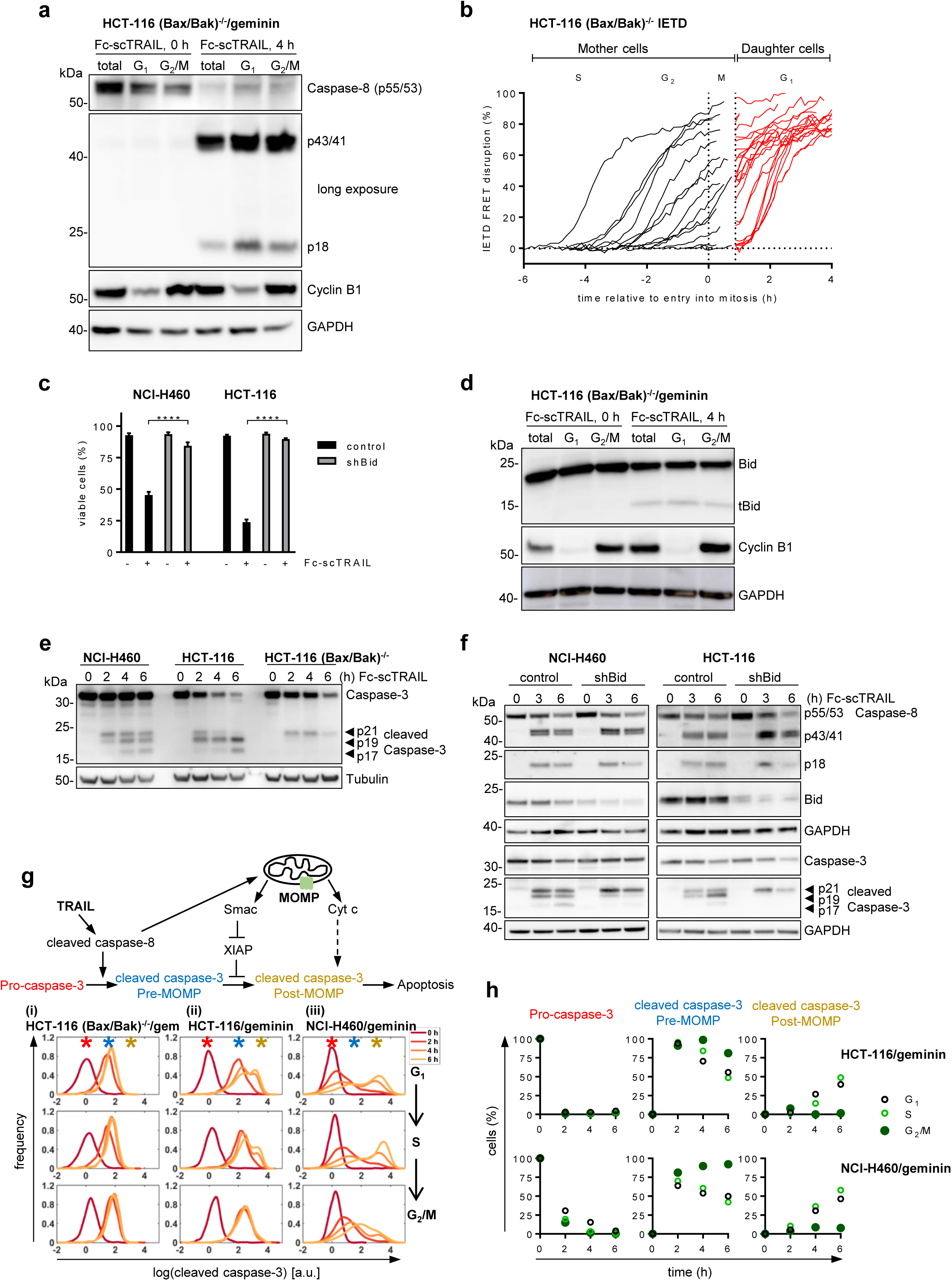
Caspase-8 remains active throughout mitosis, but efficient caspase-3 activation is suppressed during G_2_/M phases. **a** Cells were treated with Fc-scTRAIL (0.6 nM) as indicated, and G_1_ and G_2_/M subpopulations were separated by FACS. Populations were analysed by immunoblotting. Blots are representative of three independent experiments. **b** HCT-116 (Bax/Bak)^-/-^ cells expressing an IETD FRET probe were exposed to Fc-scTRAIL (0.6 nM) and monitored by time-lapse fluorescence microscopy. Traces show FRET disruption, indicative of caspase-8 activity, in individual cells while transitioning through mitosis. Data show responses of cells from n=3 independent experiments. **c** Cells were treated with Fc-scTRAIL (0.06 nM (NCI-H460) or 0.6 nM (HCT-116)) for 24 h, harvested and stained with Annexin V-GFP and PI. Cells were analysed by flow cytometry and viable fractions are shown as mean values + SD of one representative experiment from three independent repeats. ****p<0.0001, unpaired t-tests. **d** Cells were treated and sorted as in (A) and analysed by immunoblotting. Blots are representative of three independent experiments. **e** Cells were treated with Fc-scTRAIL (0.06 nM (NCI-H460) or 0.6 nM (HCT-116)). Cell lysates were analysed by immunoblotting. Blots are representative of two independent experiments. **f** Control and Bid-depleted (shBid) cells were exposed to Fc-scTRAIL (0.06 nM (NCI-H460) or 0.6 nM (HCT-116)). Cell lysates were analysed by immunoblotting. Blots shown are representative of two independent experiments. **g** Upper panel shows a schematic representation of caspase-3 processing in type II cells in response to TRAIL receptor activation. Subpanels (i-iii) show caspase-3 cleavage in the indicated cell lines between 0 – 6 h after treatment with Fc-scTRAIL (0.6 nM for HCT-116 or 0.06 nM for NCI-H460). Population distributions of one representative experiment out of three are shown. Peaks representing cells with pro-caspase-3, limited pre-MOMP cleavage of caspase-3 or full post-MOMP caspase-3 processing are highlighted by asterisks. **h** Quantification of pro-caspase-3 and cleaved caspase-3 signals in distinct cell cycle phases. A population mixture model with three subpopulations was fitted to data shown in **g**.

Taken together, we conclude that caspase-8 is activated independently of the cell cycle phase, that caspase-8 remains active throughout mitosis and that activity can be transmitted to daughter cells. Natural caspase-8 substrates Bid and procaspase-3 can be processed in all phases of the cell cycle. However, cells analysed in G_2_/M phases can prevent full caspase-3 processing, suggesting that MOMP is not triggered these subpopulations.

### Cells evading TRAIL-induced apoptosis after transitioning through mitosis suffer sublethal damage

Since cells are capable of evading MOMP and inheriting caspase-8 activity to daughter cells following TRAIL treatment, we studied if such caspase-8 activities inevitably result in apoptosis execution after mitosis. Even though we observed cells dying after mitosis (**Figure 2**), we also noted that substantial numbers of cells remained viable after long treatment times (**Figure S1b**). As expected, cells dying after mitosis lost their mitochondrial membrane potential, a surrogate marker for MOMP, just prior to displaying apoptotic phenotypes (**Figure 4a, b**). Moreover, we occasionally observed that sibling cells may experience distinct cell fates after mitosis, with mitochondrial depolarization and apoptosis observed in one cell but not in the other (**Figure 4b**). It is therefore conceivable that cells might escape apoptosis induction subsequent to caspase-8 activation. We consequently studied if caspase activities could be observed in cells not showing overt apoptotic phenotypes following TRAIL treatment and mitosis. As a reference, TRAIL treatment caused dramatic caspase-dependent FRET disruption in non-dividing, apoptosis competent NCI-H460 and HCT116 cells expressing the IETD FRET probe (**Figure 4c**). Analysing cells passing through mitosis and subsequently either dying or failing to show an apoptotic phenotype revealed notable differences. Cells dying after mitosis displayed FRET loss similar to non-dividing cells, whereas cells that escaped apoptosis cleaved significantly less FRET probe (**Figure 4d**). Importantly, FRET efficiency in apoptosis-averting cells nevertheless differed significantly from untreated control cells, indicating that caspases were activated in sublethal amounts (**Figure 4d**). Apparently, many of these cells suffer DNA damage, as detected by phosphorylation of H2A.X, a well-established marker for DNA double strand breaks (**Figure 4e, f**). Addition of QVD-OPh or siRNA-mediated silencing of caspase-activated DNAse (CAD), the major executioner of apoptotic DNA fragmentation, suppressed pH2A.X signals, demonstrating that DNA damage was caspase-dependent (**Figure 4f, g**). Overall, these experiments therefore indicate that caspase-8 activity, which can persist through mitosis, not inevitably induces cell death in daughter cells. Cells surviving TRAIL treatment, however, display overt signs of caspase-dependent DNA damage.

**Figure 4:**
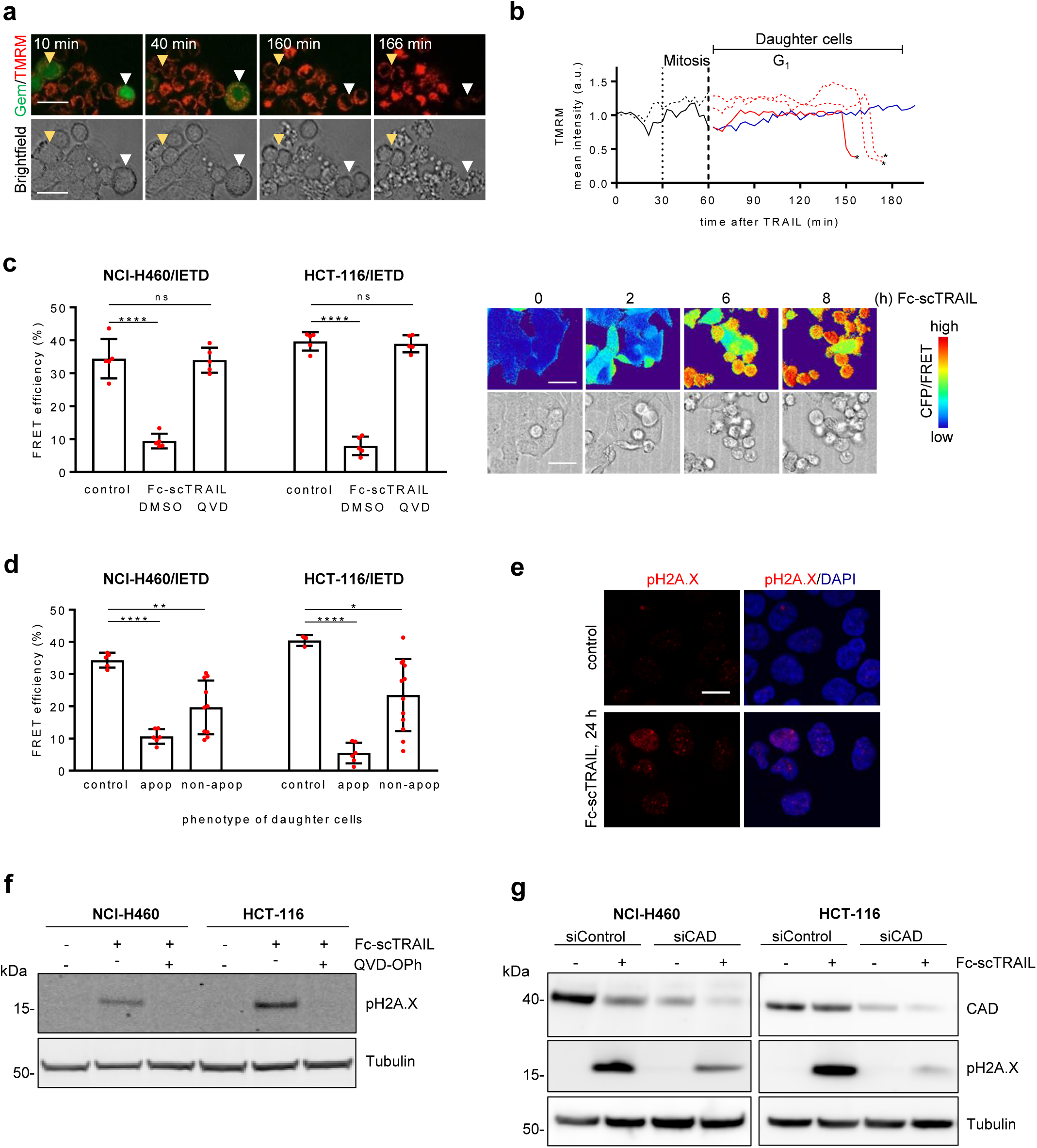
Cells evading TRAIL-induced apoptosis after transitioning through mitosis suffer sublethal damage. **a** Geminin-expressing NCI-H460 cells were loaded with TMRM (30 nM) and treated with Fc-scTRAIL (0.06 nM). White arrowhead indicates a cells progressing through mitosis, followed by mitochondrial depolarization and apoptosis of daughter cells. Yellow arrowhead indicates a cell progressing through mitosis without daughter cells committing apoptotic cell death. Scale bar = 20 µm. **b** Quantification of TMRM intensities in Fc-scTRAIL treated cells passing through mitosis. Solid and dashed lines indicate two mother cells and their corresponding daughter cells. Asterisks indicate times of apoptotic phenotypes. Blue line indicates a surviving cell. Traces are representative for n = 10 mother cells analysed. **c** NCI-H460 or HCT-116 cells expressing an IETD FRET probe were exposed to Fc-scTRAIL (0.06 nM or 0.6 nM) alone or in the presence of QVD-OPh (25 µM) and monitored by confocal microscopy. FRET efficiencies were calculated after acceptor photobleaching. Data show mean ± SD of five cells for each condition. **** p<0.0001 and ns, not significant, compared versus control (One-way ANOVA followed by Dunnett’s multiple comparison). Micrographs show FRET probe expressing HCT-116 cells treated with Fc-scTRAIL (0.6 nM), with the upper panel showing pseudo-coloured CFP/FRET emission ratios. Scale bars = 50 µm. **d** Cells were treated and monitored as in **c**. FRET efficiencies were analysed in individual cells that progressed through mitosis and for which daughter cells either showed or did not show apoptotic or non-apoptotic phenotypes. Data show mean ± SD values of control cells (n = 3-5), apoptotic cells (n = 6) and non-apoptotic cells (n = 10-12). ****p<0.0001, **p<0.01, *p<0.05. **e** NCI-H460/geminin cells were treated with Fc-scTRAIL (0.06 nM). After 24 h, adherent surviving cells were stained for pH2A.X (Ser139) and DNA and analysed by confocal microscopy. Scale bars = 10 µm. **f** Cells were treated as indicated. After 24 h, surviving cells were analysed by Western blotting. Tubulin served as loading control. **g** Control and CAD-depleted cells were treated with Fc-scTRAIL (0.3 nM) for 24 h. Cell lysates were analysed as indicated. Tubulin as loading control. Blots are representative of three independent experiments.

### Mcl-1 accumulates in cells preparing to enter mitosis

Cells delaying or escaping from TRAIL-induced apoptosis do so by averting MOMP in late stages of the cell cycle. We therefore studied which molecular mechanisms might confer this increased MOMP resistance. The anti-apoptotic Bcl-2 family members Mcl-1 and Bcl-xL, key regulators of MOMP sensitivity, have previously been reported to prevent or suppress apoptosis during extended mitotic arrest induced by anti-mitotic drugs such as paclitaxel and taxol (29–33). First, we therefore determined absolute expression quantities of both proteins in NCI-H460 and HCT-116 cells, in comparison to HeLa cells for which nM expression amounts were reported previously (34). Both cell lines expressed higher levels of Bcl-xL than Mcl-1, and NCI-H460 cells expressed approximately 9-fold higher amounts of Mcl-1 than HCT-116 cells (**Figure 5a**). The combined expression of Bcl-xL and Mcl-1 was comparable between both cell lines, though. Studying expression of both proteins across the cell cycle indicated that both cell lines substantially elevated Mcl-1 in M phase (i.e. above the expected doubling throughout the cell cycle (25, 35), as determined by flow cytometry (**Figure 5b**). This finding is in line with Mcl-1 accumulation observed across the cell cycle in synchronised U2OS cell populations (29). Higher than expected amounts of Bcl-xL were observed only in HCT-116 cells (**Figure 5b**). Immunoblotting independently confirmed the upregulation of Mcl-1 in later stages of the cell cycle (**Figure 5c**). A more detailed kinetic analysis of Bcl-xL and Mcl-1 expression across the cell cycle likewise confirmed the increase in Mcl-1, beginning in mid S phase and accumulating substantially in late G_2_/M phases (**Figure 5d**).

**Figure 5:**
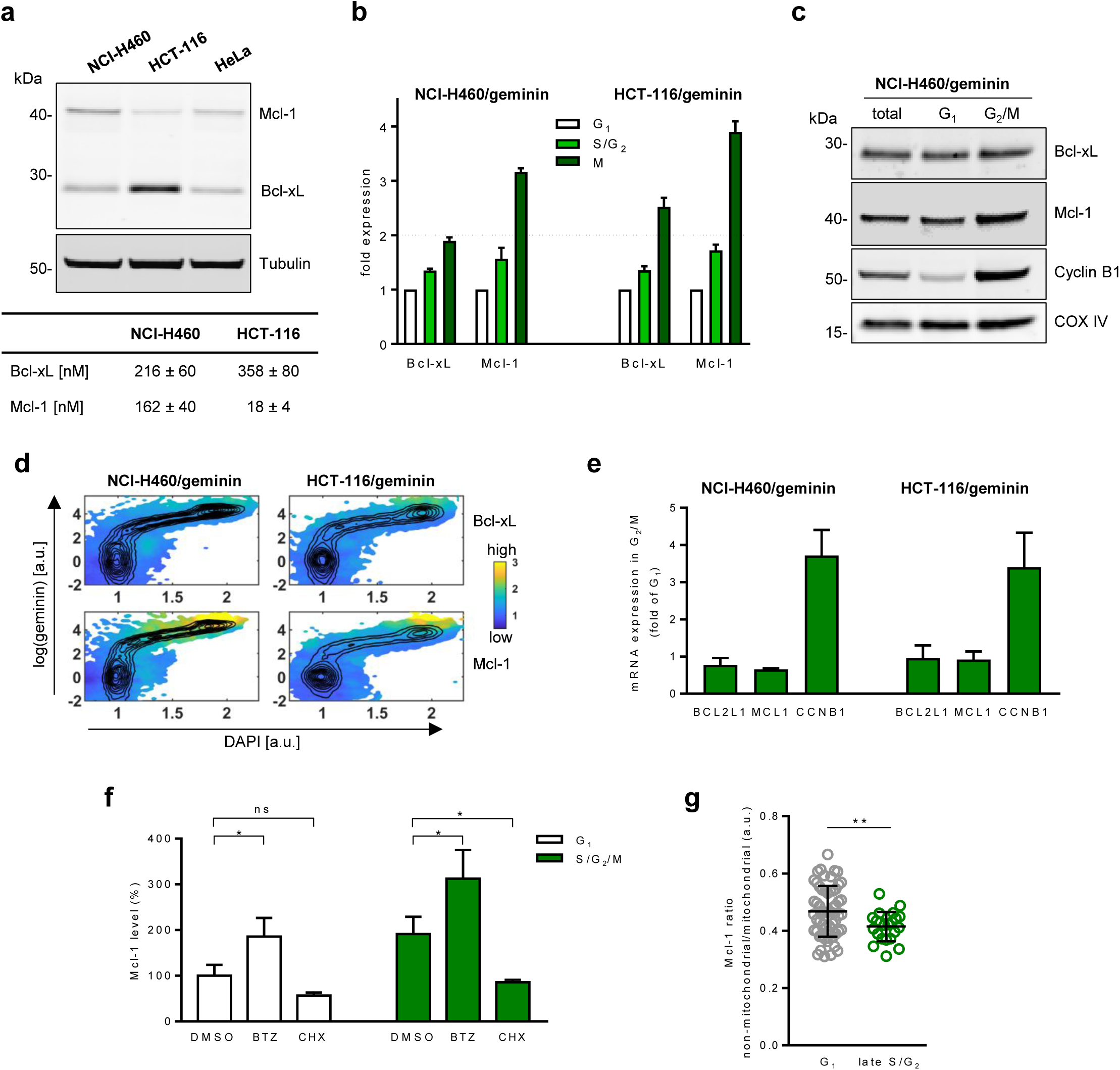
Mcl-1 accumulates in cells preparing to enter mitosis. **a** Cell lysates were immunoblotted as indicated. Tubulin served as loading control. Concentrations of Bcl-xL and Mcl-1 were quantified from signal intensities in comparison to HeLa cells, for which protein concentrations were reported (34). **b** Cells were co-stained with antibodies against Bcl-xL or Mcl-1 and VioBlue-pH3-S10, and analysed by flow cytometry. Median fluorescence intensities were normalized against signals obtained from cells in G_1_ phase. Data show fold expression (mean + SEM) from at least four independent experiments. **c** Cells were FACS-separated into G_1_ and G_2_/M populations and analysed by immunoblotting. COX IV served as loading control. Blots are representative of three experiments. **d** Cells were stained for Bcl-xL or Mcl-1 and DNA, followed by flow cytometric analysis. Representative heat maps of cell cycle-dependent expression of Bcl-xL (upper panels) and Mcl-1 (lower panels) are shown as a function of DNA content and geminin expression. **e** Cells were FACS-separated into G_1_ and G_2_/M subpopulations, followed mRNA extraction. Transcript levels of *BCL2L1* (Bcl-xL), *MCL1* (Mcl-1), and *CCNB1* (Cyclin B1) were analysed by qPCR. Data show means + SD from three independent experiments. **f** NCI-H460/geminin cells were treated with DMSO, bortezomib (BTZ, 25 µg/ml) or cycloheximide (CHX, 5 µg/ml) for 1.5 h, co-stained for Mcl-1 and DNA, and analysed by flow cytometry. Mcl-1 amounts are shown as means + SD from three independent experiments. *p<0.05 (one-way ANOVA plus Sidak’s multiple comparisons test). **g** NCI-H460/geminin cells were analysed for mitochondrial and non-mitochondrial Mcl-1 intensities. Shown are median ± SD calculated from G_1_ cells (n = 58) and S/G_2_ cells (n = 22) from two independent experiments (**p<0.01, Welch’s unequal variances t-test).

We next studied which processes might contribute to the accumulation of Mcl-1 in late cell cycle stages. In contrast to cyclin B1 mRNA, Mcl-1 mRNA did not accumulate in G_2_/M phases, so that transcriptional upregulation could be excluded (**Figure 5e**). After short periods of proteasome inhibition by bortezomib (BTZ), Mcl-1 accumulated in cells from both G_1_ and S/G_2_/M phases (**Figure 5f**). In reverse, Mcl-1 amounts decreased in both groups of cells when inhibiting protein translation by cycloheximide (CHX) (**Figure 5f**). Mcl-1 is therefore translated and degraded in early and in later stages of the cell cycle, so that a combination of alterations in both translation and/or degradation rates would contribute to a net increase in Mcl-1 amounts prior to mitosis. Multi-domain Bcl-2 family members continuously shuttle between mitochondrial membranes and the cytosol (36). Since proteasomal degradation is limited to the soluble protein fraction, increased Mcl-1 stability could also be achieved by a higher membrane association during later cell cycle stages. Studying subcellular Mcl-1 distributions indeed indicated a significant redistribution towards the mitochondrial fraction in later cell cycle stages (**Figure 5g**). Overall, these data suggest that the pronounced accumulation of Mcl-1 towards mitosis and its redistribution towards mitochondrial membranes could contribute to delaying or preventing apoptosis in cells transitioning through mitosis.

### Antagonizing Mcl-1 restores apoptosis sensitivity during cell cycle progression

To assess if anti-apoptotic Bcl-2 family members interfere with apoptosis signal transduction during later stages of the cell cycle, we next studied cells in which Bcl-2, Bcl-xL or Mcl-1 were pharmacologically inhibited. In NCI-H460 cells, Bcl-2 inhibition with ABT-199 or Bcl-xL inhibition by WEHI-539 moderately reduced t_death_ times in cells passing through mitosis, whereas Mcl-1 inhibition by S63845 was significantly more effective (**Figure 6a**). A similar shortening of t_death_ times was observed in HCT-116 cells upon Mcl-1 inhibition (**Figure 6a**). Additionally, also Bcl-xL inhibitor WEHI-539 prevented prolonged t_death_ times in HCT-116 cells passing through mitosis (**Figure 6a**), possibly due to their high dependency on Bcl-xL expression (**Figure 5a**). Since Mcl-1 inhibition averts delays in apoptosis signalling in cells preparing to enter mitosis, it would be expected that overall less cells are able to progress through mitosis under these treatment conditions. Indeed, the frequency of mitotic events observed for TRAIL-treated NCI-H460 and HCT-116 cells was significantly reduced when co-treated with S63845 (**Figure 6b**). Importantly, none of the inhibitors reduced cell viability or induced notable apoptosis in absence of TRAIL treatment (**Figure S4a, b**). Using A-1210477 as an alternative Mcl-1 inhibitor provided similar findings (**Figure S4c, d**). Similar to pharmacological Mcl-1 inhibition, depleting Mcl-1 expression by siRNA also abolished prolonged t_death_ times and reduced the frequency of cell divisions following TRAIL treatment (**Figure 6c-e**). In line with these findings, flow cytometric studies revealed that TRAIL-treated cells no longer suppress caspase-3 processing in G_2_/M phases (**Figure 6f, g**, compare to **Figure 3g, h**). Instead, the kinetics of caspase-3 processing were largely identical in cells from G_1_, S and G_2_/M phases.

**Figure 6:**
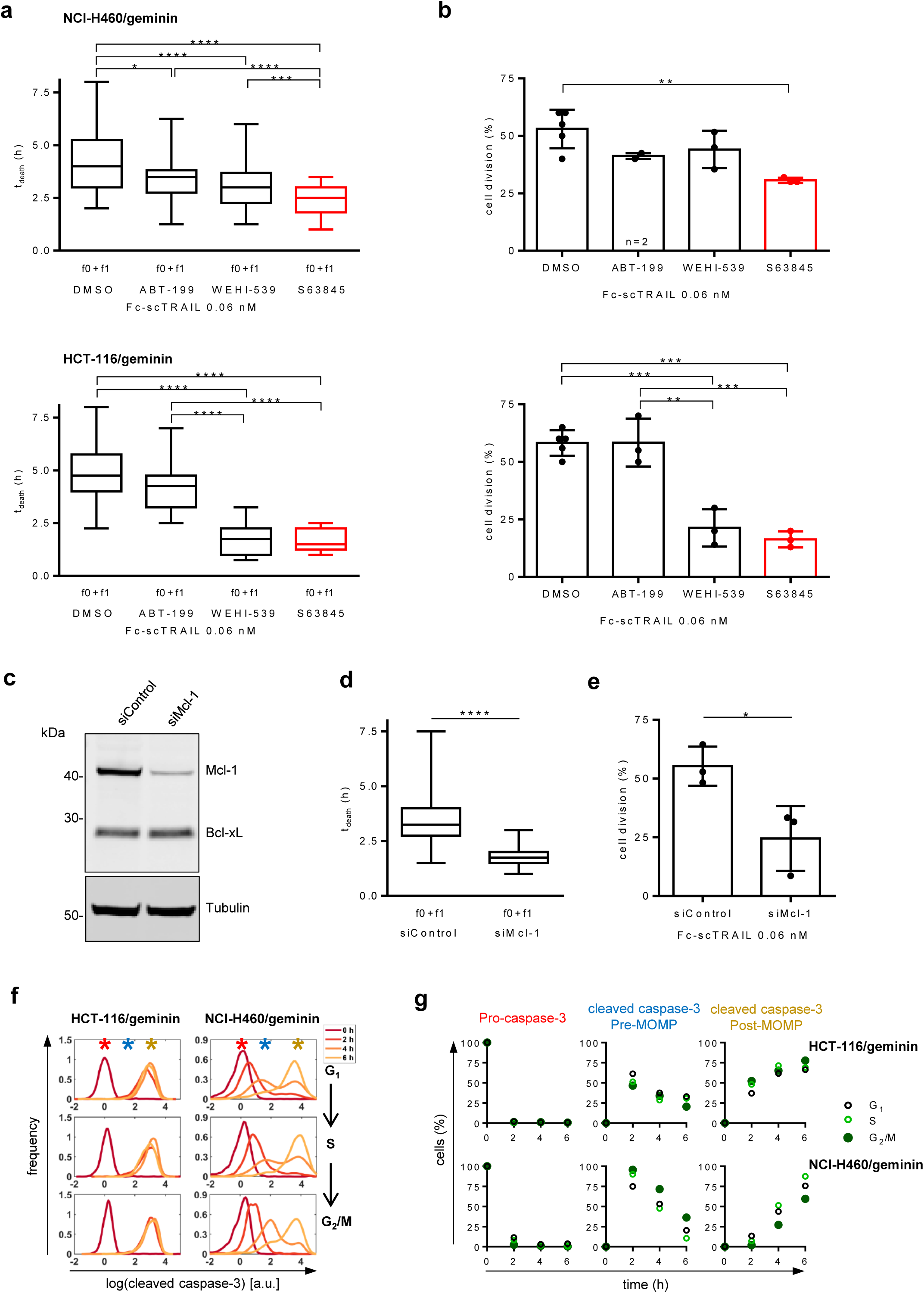
Antagonizing Mcl-1 restores apoptosis sensitivity during cell cycle progression. **a** Cells were monitored by time lapse imaging and stimulated with Fc-scTRAIL in presence or absence of Bcl-2 family antagonists (10 µM ABT-199 (Bcl-2), 10 µM WEHI-539 (Bcl-xL), 1 or 10 µM S63845 (Mcl-1). t_death_ times were determined as in Fig.2. Box and whisker plots show t_death_ for cells that die after (f0+f1) mitosis. Shown are medians with interquartile ranges, plus min to max range from n = 60-115 cells observed in two to three independent experiments. ****p<0.0001, ***p<0.01, *p<0.05 (Kruskal-Wallis test followed by Dunn’s multiple comparison test). **b** The percentage of S/G_2_/M cells that progress through mitosis upon Fc-scTRAIL exposure was determined from cells analysed in (A). Data show means ± SD (***p<0.001, **p<0.01; one-way ANOVA with Tukey’s multiple comparison test). **c** NCI-H460/geminin cells were transfected for 24 h and the amount of Bcl-xL or Mcl-1 was determined by immunoblotting. Tubulin served as loading control. **d** Death times after depletion of Mcl-1. NCI-H460/geminin cells were treated with Fc-scTRAIL (0.06 nM). Box and whisker plots show time to death (t_death_) for cells that die after mitosis (f0+f1). Shown are medians with interquartile ranges, plus min to max ranges calculated from at least 75 cells from three independent experiments. **** p<0.0001 (one-way ANOVA with Tukey’s multiple comparison test). **e** The percentage of S/G_2_/M cells undergoing mitosis was calculated from experiments analysed in (D). Data are means ± SD from n = 3 experiments. *p<0.05, unpaired t-test. **f** As in Figure 3g, panels show caspase-3 cleavage between 0 – 6 h after treatment with Fc-scTRAIL (0.6 nM for HCT-116 or 0.06 nM for NCI-H460) in presence of Mcl-1 inhibitor A-1201477 (10 µM). Peaks representing cells with pro-caspase-3, limited pre-MOMP cleavage of caspase-3 or full post-MOMP caspase-3 processing are highlighted by asterisks. **g** Quantification of pro-caspase-3 and cleaved caspase-3 signals in distinct cell cycle phases. A population mixture model with three subpopulations was fitted to time course data shown in **f**. Note the absence of kinetic differences between the subpopulations (compare also to Figure 3h).

Together, these findings demonstrate that Mcl-1 confers transient resistance to TRAIL-induced extrinsic apoptosis in cells entering and passing through mitosis, and that this resistance can be antagonized by inhibiting or depleting Mcl-1. Consequently, the combination of TRAIL and Mcl-1 inhibition completely abrogated long term survival of NCI-H460 cells while HCT-116 cell survival was markedly reduced by co-treatment with Bcl-xL or Mcl-1 inhibitors (**Figure S4e, f**).

## Discussion

Here, we showed that cells progressing towards and passing through mitosis delay TRAIL-induced extrinsic apoptosis. Importantly, cells transition through mitosis with active caspase-8, are protected from mitochondrial apoptosis, but nevertheless accumulate caspase-dependent sublethal DNA damage. Upregulation of Mcl-1 and, to a lower extent, Bcl-xL confer transmitotic apoptosis resistance, with BH3-mimetics targeting these proteins preventing unwanted escape from extrinsic apoptosis.

To obtain insight into the relationship between extrinsic apoptosis signalling and cell cycle progression, we employed fluorescence-based time lapse imaging of non-synchronized cell populations. This circumvented the limited temporal resolution and accuracy of traditional chemical synchronization (17). Furthermore, cell cycle arrest for synchronization purposes not only would have interfered with normal cell cycle progression but could have further confounded the analyses by inducing stress responses that modify apoptosis sensitivities in distinct cell cycle phases.

Cells delay or prevent apoptosis execution when proceeding towards cell division, with the decision on whether to favour mitosis over apoptosis being made approximately half way through S phase. Decision making therefore substantially precedes entry into mitosis and gives rise to prolonged periods of increased apoptosis resistance, explaining also why cell death events are more frequently observed during G_1_ phase. The cell-to-cell signalling heterogeneity during the pre-MOMP phase so far was solely attributed to protein expression noise across cell populations (19), while here we show that cell cycle progression adds an additional and significant non-random layer of cell death regulation. Since we observed similar delays in extrinsic apoptosis signalling when stimulating NCI-H460 cells with FasL, transmitotic apoptosis resistance seems independent of the choice of death ligand.

Mcl-1 is the primary factor conferring transmitotic resistance to extrinsic apoptosis. A key role for Mcl-1 in delaying or suppressing cell death has been reported previously at conditions of mitotic arrest (29,32,33,37,38), and various E3 ligases have been reported that could contribute to eliminating Mcl-1 (39). In our setting, where mitosis proceeds unperturbed, cells rapidly deplete Mcl-1 upon entry into G_1_ phase, with many daughter cells regaining the competency to die due to their inherited caspase-8 activity. If one or several of the known Mcl-1 E3 ligases is implicated in rapid Mcl-1 depletion upon regular exit from mitosis remains to be studied.

We found that Mcl-1 steadily accumulates from mid S phase on, matching trends reported for synchronized cell populations (29), but also to additionally sharply increase at the very end of the cell cycle. Apoptosis resistance must be very high during this period, since we never observed apoptosis execution during mitosis itself, despite analysing hundreds of cells. Since transcription is largely blocked upon entry into mitosis, Mcl-1 accumulation could arise from continued translation during M phase, possibly combined with suppressed degradation, continuing trends we observed when analysing the pooled fraction of cells in G_2_/M phases. Indeed, Mcl-1 mRNA continues to be translated during mitosis, as shown in other studies (33, 40). Besides translation and proteasomal degradation contributing to defining Mcl-1 amounts throughout the cell cycle, we observed a significant redistribution of Mcl-1 towards mitochondrial membranes in later cell cycle phases. This offers a currently possibly underappreciated mechanism by which Mcl-1 could be depleted from the degradable soluble fractions and which thereby could contribute to protein stabilisation. Bcl-xL likewise accumulates across the cell cycle but lacks the sharp increase in its amounts during mitosis. While Bcl-xL can reduce apoptosis sensitivity during mitotic arrest by paclitaxel in breast cancer cells (31), in our cell line models Mcl-1 inhibition more consistently avoided transmitotic apoptosis resistance. Mcl-1 inhibition likewise outperforms Bcl-xL inhibition in apoptosis induction by mitotic driver drugs, such as aurora kinase inhibitors (41). The third major anti-apoptotic Bcl-2 family member, Bcl-2 itself, is not regulated by cell cycle progression (29) and instead apparently is inactivated by phosphorylation prior to/upon entry into mitosis (42). Similar activity-suppressing phosphorylation has also been reported for Bcl-xL (31,42,43). Overall, this indicates that in particular during mitosis, apoptosis resistance would be expected to strongly rely on Mcl-1. Earlier studies reported the suppression of caspase-8 activation by Ser-387 phosphorylation in cells arrested in mitosis (44, 45), whereas we observed that caspase-8 can be activated and remains active in all phases of the cell cycle. The phosphorylation dependent inhibition of caspase-8 activation therefore might be restricted to cells arrested in mitosis rather than manifesting prominently during regular cell cycle progression or represent a layer of regulation that does not prominently display in the cellular model systems studied here. Likewise, additional regulatory steps downstream of MOMP, such as the inactivation of caspase-9 by phosphorylation during mitosis (46), could further contribute to establishing transiently higher apoptosis resistance.

A more general question arises regarding why cells would need to raise their apoptosis resistance in preparation for and during mitosis. During S phase, cells expend substantial amounts of energy on synthesis and growth processes, which in themselves might impose increased basal stress against which cells protect themselves. Alternatively or in addition, proliferation might be associated with mild or moderate intrinsic pro-apoptotic activities that are not supposed to lead to cell death. Indeed, various apoptotic proteins have been ascribed roles independent of cell death. For example, cellular remodelling during development, cell fate determination of stem cells, as well as in the regulation of cytoskeletal dynamics partially depend on non-apoptotic caspase signalling (47). Interestingly, RIPK1-containing complexes form during mitosis to activate caspase-8, which contributes to orderly chromosome segregation, without these cells showing signs of apoptosis (48). Furthermore, caspase-8 activity seems required to overcome the G_2_/M checkpoint by p53 destabilization (49). This would imply that apoptosis thresholds must be elevated during mitosis to withstand unwanted apoptosis. In line with this, cells in which Mcl-1 and Bcl-xL are pharmacologically inhibited spontaneously and preferentially die during mitosis (41). Evasion from extrinsic apoptosis thus would arise as an unwanted side effect, which however could have substantial implications for the efficacy by which the immune system can clear transformed cells and, by extension, the risk to relapse from immune stimulatory or death ligand-based therapy (5,50,51). Inhibitors of Mcl-1 and Bcl-xL might therefore have a space as co-treatments in such scenarios, especially since Mcl-1 is highly amplified in various cancers (52). Moreover, the Mcl-1 dependency of malignant neoplasms generally might be elevated, since recent CRISPR-Cas9 screening identified Mcl-1 and, with a lower score, Bcl-xL as attractive priority targets in colorectal cancers (53).

We noted caspase-dependent DNA damage in cells escaping TRAIL induced apoptosis by passing through mitosis. DNA damage in TRAIL-surviving cells was previously described (54), and such damage could arise from moderate activation of caspase-3 either through direct cleavage by caspase-8 and/or the inefficient activation of MOMP, resulting in minority MOMP, a process of inefficient MOMP induction that can contribute to further cell transformation in response to Bcl-2 family antagonism and to inflammatory cytokine production in infection scenarios (55, 56). Since caspase-8 can also cleavage DEVD sequences, such as found in PARP (26, 57), it cannot be excluded that caspase-8 itself likewise contributes to DNA damage.

Taken together, our study shows that regular cell cycle progression modulates signal transduction during extrinsic apoptosis, with Mcl-1 modulating cellular decision making between death, proliferation and survival from inefficient apoptosis. These findings not only extend our understanding of the temporal regulation of extrinsic apoptosis susceptibility but also are relevant for better understanding cellular mechanisms contributing to escape from apoptosis induction.

## Materials and methods

### Maintenance of cell lines

Cells were cultured in RPMI 1640 medium supplemented with 10% fetal bovine serum (FBS) and kept at 37°C in 5% CO_2_ humidified atmosphere. HCT-116 cells were obtained from Interlab Cell Line Collection (Italy). NCI-H460, HeLa and HT1080 cells were obtained from ATCC (LGC Standards GmbH, Germany). Cell isolates from melanoma metastases were kindly provided by D. Kulms (Department of Dermatology, University of Dresden, Germany) and obtained as part of routine resections at University Hospital Dresden, under the auspices of the local Ethics Committee (ethical approval number EK335082018). Platinum-E (Plat-E) cells were kindly provided by M. Olayioye, University of Stuttgart, Germany. Cell lines were routinely tested by multiplex human cell line authentication test (Multiplexion GmbH, Germany). NCI-H460/geminin and NCI-H460/Bcl-2 cells were described previously (7, 22), HCT-116 (Bax/Bak)^-/-^ cells were kindly provided by R. Youle (National Institutes of Health, USA).

### Antibodies and reagents

Primary antibodies were as follows: mouse anti-β-Actin (8H10D10, Cell Signaling), rabbit anti-Bak (Cell Signaling), rabbit anti-Bax (Cell Signaling), rabbit anti-Bcl-xL (Cell Signaling), rabbit anti-Bid (Cell Signaling), mouse anti-Caspase-8 (1C12, Cell Signaling), rabbit anti-Caspase-3 (Cell Signaling), rabbit anti-COX IV (3E11, Cell Signaling), rabbit anti-Cyclin B1 (D5C10, Cell Signaling), mouse anti-GAPDH (D4C6R, Cell Signaling), rabbit anti-Mcl-1 (Cell Signaling), mouse anti-PARP (BD Biosciences), rabbit anti-Phospho-Histone H2A.X (Ser139) (20E3, Cell Signaling), rabbit anti-Phospho-Histone H3 (Ser10) Pacific Blue™ Conjugate (D2C8, Cell Signaling), mouse anti-α-Tubulin (DM1A, Cell Signaling), rabbit anti-cleaved caspase-3 AF647 Conjugate (Asp175) (D3E9, Cell Signaling), mouse anti-TRAIL-R1 (MAB347, R&D Systems), mouse anti-TRAIL-R2 (MAB6311, R&D Systems), mouse IgG1 κ isotype control (BD Biosciences), mouse IgG2b κ isotype control (BD Biosciences), mouse IgG2a κ isotype control (BD Biosciences), mouse anti-FLAG-PE conjugate (BioLegend), rabbit (DA1E) mAb IgG XP® isotype control (Cell Signaling), rabbit IgG isotype control AF647 Conjugate (Cell Signaling). Secondary antibodies were goat anti-rabbit IgG (H+L) secondary antibody, Alexa Fluor 647 (Thermo Fisher Scientific), AF647-conjugated goat anti-mouse IgG+IgM (Dianova), IRDye®680RD goat anti-rabbit/mouse IgG (LI-COR Biosciences), IRDye®800CW goat-anti-rabbit/mouse IgG (LI-COR Biosciences) and peroxidase-conjugated goat anti-rabbit/mouse IgG (Dianova).

Fc-scTRAIL was produced as described previously (23), FasL-Fc was kindly provided by H. Wajant (University Hospital Würzburg, Germany) and Annexin V-GFP was produced in-house. DMSO was purchased from Carl Roth, ABT-199 was from Active Biochem, A-1210477 and Q-VD-Oph from Selleckchem, S63845 and WEHI-539 from APExBIO and bortezomib (PS-341) from UBPBio. TMRM, MitoTracker™ Red CMXRos and DAPI were obtained from Invitrogen, propidium iodide (PI) and cycloheximide (CHX) were purchased from Sigma. Puromycin and G-418 were obtained from AppliChem GmbH.

### Cell death measurements

Cells were treated as indicated and viability was determined by crystal violet staining after 24 h of treatment. Absorbance was measured in a micro plate reader at λ = 550 nm and normalized to those of control cells. Alternatively, cells were harvested, stained with Annexin V-GFP/PI and viability was analysed by flow cytometry (MACSQuant, Miltenyi Biotec, Germany). To analyse the long-term survival, cells were treated as indicated for 24 h. Cells were washed to remove dead cells and surviving cells were cultivated in fresh medium for 7 days. Viability was then determined by crystal violet staining.

### Plasmids and transfections

Plasmid pEFpuro-hGem(1/110) was obtained by subcloning cDNA encoding mAG-hGem(1/110) (21) into pEFpuro. The plasmid pSCAT8 encoding the caspase-8 FRET probe CFP-IETD-Venus was described previously (27), and a SCAT8 variant in a pIRESpuro3 backbone was obtained by subcloning the CFP-IETD-Venus sequence. Cells were transfected using Lipofectamine®LTX (Thermo Fisher Scientific) according to the manufacturer’s protocol. 24 h post transfection, cells were cultured in medium supplemented with appropriate selection antibiotics and fluorescent clones were isolated or enriched after two weeks.

The pQCXIPN/ecoR plasmid encoding the receptor for ecotropic murine retroviruses was a kind gift of M. Olayioye, University Stuttgart, Germany (58). Cells were transfected using Lipofectamine®LTX. 24 h post transfection, cells were incubated with supernatant from Plat-E cells transfected with pSuper RV or pSUPER RV Bid (#2) (kindly provided by S. Tait, University of Glasgow, UK) (59). Cells were then selected in medium with puromycin and analyzed for Bid expression.

Silencer®Select Negative Control siRNA #2 (4390846, Thermo Fisher Scientific), Silencer®Select siRNA targeting Mcl-1 (s8583) or Silencer®Select siRNAs targeting CAD (s4060, s223407) were transfected using Lipofectamine®RNAiMAX (Thermo Fisher Scientific) according to the manufacturer’s protocol.

### Western Blotting

Cells were lysed in solubilization buffer (50 mM Tris-HCl pH 7.5, 150 mM NaCl, 1 mM EDTA, 1% (v/v) TritonX-100 plus Complete Protease Inhibitors (Roche) and 1 mM DTT), incubated on ice for 10 min and centrifuged at 13,200 rpm and 4°C. Protein concentrations were determined by Bradford assay (Bio-Rad). Proteins were separated by SDS-PAGE (4-12% NuPAGE® Novex Bis-Tris gels, Thermo Fisher Scientific) and transferred to nitrocellulose membranes (iBlot® Gel Transfer Stacks, Thermo Fisher Scientific). Membranes were then blocked with 1% blocking reagent (Roche) or 5% BSA in TBS containing 0.05% Tween-20 (TBST) and incubated with primary antibodies followed by POD-conjugated secondary antibodies, and detection was performed using Pierce ECL Western Blotting Substrate (Thermo Fisher Scientific) using an Amersham Imager 600 (GE Healthcare). Alternatively, membranes were incubated with IRDye® secondary antibodies and detection was performed using an Odyssey Imaging system (Li-COR Biosciences).

### Quantitative RT-PCR analysis

Total RNA was extracted from cells using the NucleoSpin RNA kit (Machery & Nagel). Synthesis of cDNA and amplification of *BCL2L1* (Bcl-xL), *MCL1* (Mcl-1), *CCNB1* (Cyclin B1) or *RPLP0* (60S acidic ribosomal protein P0) was performed using enzyme mix (Power SYBR™ Green RNA-to-CT™ 1-Step kit) and the following primer:

Bcl-xL-for: 5’-AGGCGGATTTGAATCTCTTTC-3’,

Bcl-xL-rev: 5’-CCCGGTTGCTCTGAGACATT-3’,

Mcl-1-for: 5’-AACTGGGGCAGGATTGTGAC-3’,

Mcl-1-rev: 5’-CAAACCCATCCCAGCCTCTT-3’,

CCNB1-for: 5’-ACCTGTGTCAGGCTTTCTCTG-3’,

CCNB1-rev: 5’-CTGACTGCTTGCTCTTCCTCA-3’,

RPLP0-for: 5’-CTCTGCATTCTCGCTTCCTGGAG-3’,

RPLP0-rev: 5’-CAGATGGATCAGCCAAGAAGG-3’.

Relative expression was calculated using the delta delta Ct method and was plotted as fold of G_1_ mRNA amounts.

### Coupled analysis of cell cycle and caspase-3 processing

Cells were harvested, fixed, permeabilised and incubated with Alexa Fluor 647- labelled caspase-3 antibody or isotype control antibody for 60 min, followed by DNA staining (1 µg/ml DAPI) for 10 min. Cells were then washed and analysed by flow cytometry (MACSQuant). Cleaved caspase-3 subpopulations in distinct cell cycle phases were quantified by multi-experiment mixture model analysis based on three lognormal distributions, using the MATLAB Toolbox MEMO (60). Mixture model parameters, comprising subpopulation mean and variance, and subpopulation weights for all time points and cell cycle stages were derived by minimizing the negative logarithm of the likelihood to observe the experimental data.

### Live-cell imaging, immunostaining and image analysis

For live cell imaging, cells were plated on 35 mm glass-bottom dishes (CellView Cell Culture Dish, Greiner Bio One) in phenol red free RPMI 1640 containing 10% FBS. Images were acquired for 20 h in 15 min intervals at 37°C and 5% CO_2_ on a Zeiss Cell Observer microscope equipped with an Axiocam MRm CCD camera, a Plan-Apochromat 20x/0.8 objective. Fluorescence of mAG-hGeminin(1/110) was acquired with a 470 nm LED module combined with a 62 HE filter set (Carl Zeiss). Then, medium containing Fc-scTRAIL alone or in combination with DMSO or the indicated inhibitors was added and cells were imaged for another 10 h. Randomly chosen cells were tracked and the time of division, onset of geminin expression, subsequent division and t_death_ of cells were recorded. Cells were defined as apoptotic upon onset of membrane blebbing.

FRET-based imaging was performed on a Zeiss Cell Observer SD Spinning Disk microscope, equipped with an Axiocam 503 mono CCD camera. For ratiometric FRET imaging, CFP and FRET channels were acquired with a 445 nm diode excitation laser combined with 485/30 nm (CFP) and 562/45 nm (FRET) emission filters. Venus was excited with a 514 nm laser combined with a 562/45 nm emission filter. Images were acquired in 2×2 binning mode at 37°C and 5% CO_2_ every 15 min for 20 h. Cells were then treated with Fc-scTRAIL and imaged for another 8-10 h. Subsequently, Venus was bleached in the final frame with a 515 nm bleaching laser using a UGA42-Firefly illumination system (Rapp Optoelectronics, Germany). The CFP/YFP emission ratio upon background subtraction over time, the FRET efficiency of the final frame and the FRET probe cleavage were calculated using Zen blue 2.3 software (Zeiss). To record mitochondrial membrane potentials, cells were stained with 30 nM TMRM for 30 min at 37°C and imaged (561 nm excitation laser combined with a 600/50 nm emission filter) in 3 min intervals for 6 h post treatment. Image analysis was performed with Zen Blue 2.3 software (Zeiss). Cell divisions and the time from Fc-scTRAIL addition to apoptotic cell death were analysed in parallel. For pH2A.X staining, cells grown on coverslips were treated for 24 h, dead cells were removed by washing, and surviving cells were fixed, permeabilised and immunostained for pH2A.X followed by DAPI-based nuclear staining. Images were acquired on a Zeiss Axio Observer SD Spinning Disk microscope using 405, 488, and 638 nm excitation with a PlanApochromat 63x/1.4 oil objective and an Axiocam 503 Mono CCD camera. Images were processed with the ZEN software (Zeiss).

To analyse Mcl-1 subcellular localization, NCI-H460/geminin cells grown on coverslips were incubated with MitoTracker Red CMXRos (100 nM) for 30 min before fixation, permeabilisation and immunostaining for Mcl-1. Nuclei were counterstained using DAPI. Images were acquired on a Zeiss Axio Observer SD Spinning Disk microscope equipped with a PlanApochromat 40x/1.4 oil objective and an Axiocam 503 Mono CCD camera. Geminin was excited with a 488 nm diode laser using a 525/50 nm emission filter, the AF647 dye was excited with a 638 nm diode laser using the 690/50 nm emission filter and DAPI was excited with a 405 nm diode laser using the 450/50 nm emission filter. MitoTracker Red was excited with a 561 nm diode laser using a 575/50 nm emission filter. Maximum intensity projections of the acquired z-stacks were calculated and used for analysis with Cell Profiler 3.1.9 (61). In brief, masks for nuclei, mitochondria and cytoplasm (cell mask excluding mitochondria) were segmented, mean fluorescence intensities were measured and the ratio non-mitochondria/mitochondria was plotted for geminin-negative (G_1_) or geminin-positive (late S/G_2_) cells.

### Mathematical modelling

The mathematical model serving as a reference for independent progression of apoptosis and cell cycle signalling was implemented in MatLab (The MathWorks, UK). Detailed information on model development and parameterization are reported elsewhere (24).

### Flow cytometry

For intracellular protein staining, cells were fixed with 4% paraformaldehyde, washed with PBS supplemented with 3% (w/v) FBS, and permeabilised with FACS^TM^ permeabilizing solution 2 (BD Biosciences, Germany). Cells were incubated with primary antibodies diluted in PBS/3% FBS for 60 min on ice followed by two washing steps. Cells were then incubated with Alexa Fluor 647-conjugated secondary antibody diluted in PBS/3% FBS. After washing, cells were incubated with phospho-Histone H3 (Ser10) Pacific Blue™ for 30 min and subsequently analysed by flow cytometry. To analyse the effect of bortezomib (BTZ) or cycloheximide (CHX) on Mcl-1 levels, cells were treated with DMSO, BTZ (25 µg/ml) or CHX (5 µg/ml) for 1.5 h before fixation. For staining of surface TRAIL-R1 and TRAIL-R2, cells were resuspended in PBA containing primary antibody or corresponding isotype control and incubated for 45 min on ice. After washing, cells were incubated with AF647- conjugated secondary antibody in PBA for 45 min on ice. Cells were fixed, permeabilised and incubated with phospho-Histone H3 (Ser10) Pacific Blue™ for 30 min, washed again and analysed by flow cytometry (MACSQuant). To analyse the binding of Fc-scTRAIL, cells were incubated with 0.06 nM Fc-scTRAIL for 15 min, washed and incubated with anti-FLAG-PE conjugate on ice for 30 min. Cells were washed and analysed by flow cytometry.

### Preparative cell sorting by FACS

To analyse protein amounts in G_1_ (geminin-negative) or G_2_/M (representing the highest 20% of geminin-positive cells) phases, cells were harvested, pelleted and washed with PBS supplemented with 10% FBS. Cells (2×10^6^/ml) were then separated according their mAG-geminin intensity (excitation 494 nm/emission 519 nm) using a FACSAria™III cell sorter (BD Biosciences) at 4°C. Sorted subpopulations were centrifuged and analysed by Western blotting or used for RNA isolation and quantitative RT-PCR analysis.

### Statistical Analysis

Statistical analysis was performed using PRISM 7.05 (GraphPad Software).

## Acknowledgements

The authors acknowledge Drs. D. Kulms (University of Dresden), M. Olayioye (University of Stuttgart), S. Tait (University of Glasgow), H. Wajant (University of Würzburg) and R. Youle (NIH Bethesda) for providing crucial materials. The authors thank Beate Budai and Nathalie Peters for expert technical assistance.

## Author contributions

Study design: NP, MR; Experimental work: NP, AL, JSF, IH, JS, DS; Data analysis: NP, AL, DI, KK, JSF, IH, JS, SE, DS; Contributing resources and expertise: FA, PS, MR; Manuscript writing: NP, MR.

## Supplemental Figure Legends

**Figure S1: Apoptotic cell death in response to Fc-scTRAIL.**

**a** Schematic representation of mAG-hGeminin(1/110) expression across the cell cycle.

**b** Cells were treated with Fc-scTRAIL for 24 h and stained with crystal violet. Data show mean values ± SD from a representative experiment. Similar results were obtained in two independent repeats. Nonlinear regression using a sigmoid dose-response fit was performed in GraphPad prism 7.05.

**c** Cells were treated with Fc-scTRAIL (0.06 nM for NCI-H460; 0.6 nM for HCT-116) and QVD-OPh (25 µM) as indicated. Whole cell extracts were analysed for cleaved caspase-8, cleaved caspase-3 and cleaved PARP. COX IV served as loading control.

**d** Cells were treated with Fc-scTRAIL (0.06 nM) and QVD-OPh (25 µM) for 24 h, harvested and stained with Annexin V-GFP and PI. Cells were analysed by flow cytometry and the viable fractions (Annexin V-GFP and PI negative) are shown as mean + SD from three independent experiments.

**e** Representative time-lapse images of HCT-116/geminin cells treated with Fc-scTRAIL (0.06 nM). Scale bar represents 50 µm. Cells dying prior to division are highlighted by black arrowheads. Red arrowhead tracks a cell dying after division.

**Figure S2: TRAIL death receptor amounts and Fc-scTRAIL binding expectedly double across the cell cycle.**

**a and b** Times required to die (t_death_) of HeLa (A) or HT1080 (B) cells were plotted following treatment with Fc-scTRAIL (0.06 nM for HeLa, 0.6 nM for HT1080). Plots show medians, 2^nd^ and 3^rd^ quartiles from 60 cells from two independent experiments (**** p<0.0001 Mann-Whitney U test).

**c** NCI-H460/geminin cells were treated with FasL-Fc (0.5 nM) and t_death_ values were plotted from 60 cells from two independent experiments. Plots show medians, 2^nd^ and 3^rd^ quartiles (**** p<0.0001 Mann-Whitney U test).

**d** Death times of melanoma cells isolated from metastases in response to treatment with Fc-scTRAIL (0.06 nM). Plots show medians, 2^nd^ and 3^rd^ quartiles from at least 50 cells from two independent experiments (**** p<0.0001 Mann-Whitney U test).

**e** Scatter plot of geminin versus pH3-Ser10 intensity used to define G_1_ (grey), S/G_2_ (green) and M (dark green) subpopulations of HCT-116/geminin cells.

**f** Cells were stained for cell surface TRAIL-R1 or TRAIL-R2. Cells were then fixed, permeabilised, counterstained for pH3-Ser10 and analysed by flow cytometry. The median fluorescence intensity values of cells in S/G_2_ (geminin-positive) and M (geminin/pH3-positive) phases were normalized to the respective values in G_1_ (geminin-negative) cells. Shown are mean expression + SEM from three independent experiments.

**g** Scatter plot of a geminin and DAPI-stained control HCT-116/geminin cell population used to define G_1_ (grey), S (green) or G_2_/M (dark green) subpopulations.

**h** Cell cycle-associated binding of Fc-scTRAIL to TRAIL-Rs. Cells were incubated with Fc-scTRAIL for 15 min on ice, stained with PE-conjugated anti-FLAG antibody, then fixed, permeabilised, and stained with DAPI. Cells were analysed by flow cytometry. Background-corrected mean fluorescence intensities (MFI) of the PE signal in G_1_, S and G_2_/M cells from three independent experiments are shown as mean + SD.

**i** Lysates of HCT-116/geminin (wt) and HCT-116 (Bax/Bak)^-/-^/geminin ((Bax/Bak)^-/-^) were immunoblotted for Bax and Bak. Actin served as loading control. Blots are representative of two experiments.

**Figure S3:**

**a** Lysates from NCI-H460 and NCI-H460/Bcl-2 cells were immunoblotted for Bcl-2 and tubulin as loading control. Note that the tagged Bcl-2 protein (asterisk) appears at a somewhat lower molecular weight range as compared to wild-type Bcl-2.

**b** NCI-H460/Bcl-2 cells expressing an IETD FRET probe were treated with Fc-scTRAIL (0.06 nM) and monitored by time-lapse fluorescence microscopy. Traces show FRET disruption in individual cells while transitioning through mitosis. Data show responses of cells from n = 10 mother cells and respective daughter cells from three independent experiments.

**Figure S4: Interference with Mcl-1 activity suppresses the mitotic delay in apoptosis execution and abrogates long term survival of TRAIL-treated cells.**

**a** Cells were treated with the indicated concentrations of BH3 mimetics for 24 h and then stained with crystal violet. Results from one representative experiment out of three are shown. Data are mean + SD from triplicates.

**b** Cells were treated with BH3 mimetics (ABT-199, WEHI-539 (each 10 µM) or S63845 (1 µM)) and cell extracts were analysed for caspase-3 and PARP processing by immunoblotting. COX IV served as loading control. Blots are representative of two experiments.

**c** Death times after Mcl-1 antagonism with A-1210477. Cells were monitored and treated with Fc-scTRAIL (0.06 nM) as described in Figure 2, with A-1210477 added at 10 µM where indicated. Box and whisker plots show time to death (t_death_) for cells that die after (f0+f1) mitosis. Shown are medians with interquartile ranges, plus min to max ranges calculated from at least 50 cells from three independent experiments. Statistical analysis was performed using Mann-Whitney U test, **** p<0.0001.

**d** The percentage of S/G_2_/M cells undergoing mitosis was calculated from experiments analysed in (C). Data are means ± SD from n = 3 experiments. Statistical analysis was performed using unpaired t-test, **p<0.01 (NCI-H460/geminin), **p<0.01 (HCT-116/geminin).

**e and f** Cells were treated with Fc-scTRAIL (0.06 nM) and QVD-OPh (25 µM) or the indicated BH3 mimetic (10 µM) for 24 h. Surviving cells were cultivated in fresh medium for another 7 days and stained with crystal violet. One of two independent experiments is shown.

## Supplemental Movie

**Movie S1: Long-term imaging of Fc-scTRAIL-treated NCI-H460/geminin cells.** Asynchronously growing NCI-H460/geminin cells were pre-monitored for 20 h, treated with Fc-scTRAIL (0.06 nM) and imaged for 8 h, related to Figure 1. Time point 00:16 represents addition of Fc-scTRAIL.

